# Effects of habitat management on rodent diversity, abundance, and virus infection dynamics

**DOI:** 10.1101/2022.06.11.495742

**Authors:** Nathaniel Mull, Amy Schexnayder, Abigail Stolt, Tarja Sironen, Kristian M. Forbes

## Abstract

1. Biodiversity is necessary for healthy ecosystem functioning. As anthropogenic factors continue to degrade natural areas, habitat management is needed to restore and maintain biodiversity. However, the impacts of different habitat management regimes on ecosystems have largely focused on vegetation analyses, with limited evaluation of downstream effects on wildlife.
2. We compared the effects of prairie management regimes (controlled burning, cutting/haying, or no active management) on rodent communities and the viruses they hosted. Rodents were trapped in 13 existing prairie sites in Northwest Arkansas, USA during 2020 and 2021. Rodent blood samples were screened for antibodies against three common rodent-borne virus groups: orthohantaviruses, arenaviruses, and orthopoxviruses.
3. We captured 616 rodents across 5953 trap nights. Burned and unmanaged sites had similarly high abundance and diversity (H), but burned sites had a greater proportion of grassland specialists than control sites; cut sites had the highest proportion of grassland specialist species but the lowest rodent abundance and diversity.
4. A total of 38 rodents were seropositive for one of the three virus groups (34 orthohantavirus, three arenavirus, and one orthopoxvirus). Thirty-six seropositive individuals were found in burned sites, and two orthohantavirus-seropositive individuals were found in cut sites. Cotton rats and prairie voles, two grassland specialists, accounted for 97% of the rodents seropositive for orthohantavirus, and heavier individuals were more likely to be seropositive than lighter individuals.
5. *Policy implications*: Our study indicates that controlled burns lead to a diverse and abundant community of grassland rodent species when compared to other management regimes; as keystone taxa, these results also have important implications for many other species in food webs. Higher prevalence of antibodies against rodent-borne viruses in burned prairies shows an unexpected consequence likely resulting from these community structures. Ultimately, these results provide empirical evidence that can inform prairie grassland restoration and ongoing management strategies.

## Introduction

Healthy ecosystem functioning is usually dependent on biodiversity, including species, genetic, and even parasite diversity (Cardinale et al., 2006; Winder & Shamoun, 2006; Hughes et al., 2008; Duffy, 2009). However, biodiversity continues to be negatively impacted by a variety of phenomena, including climate change, pollution, and most considerably, changes in land cover and land use (Sala et al., 2000; Haines-Young, 2009; Young et al., 2010). This is exemplified by grasslands in the United States, where approximately 70% of total historical prairie habitat and 90% of tallgrass prairie habitat has been lost (Samson et al., 2004). Habitat management is a key component of efforts to restore and maintain grassland biodiversity, but outcomes vary depending on management strategies employed (Haddock et al., 2007; Turner II et al., 2007; Haines-Young, 2009).

In light of the need for large-scale restoration and protection in grassland habitats (Gerla et al., 2012), research on the effects of different management regimes on grassland vegetation is accumulating (e.g., Newbold et al., 2020; Feher et al., 2021). Quantifying changes in vegetation provides valuable insight into the benefits of different management regimes on habitat quality. However, there has been little research on the down-stream effects of grassland management practices on animal communities, and most available studies have focused on livestock (Paudel et al., 2021) or the integration of wildlife habitat into agricultural systems (e.g., Burkhalter, 2013; Lukens et al., 2020). Given that a key objective of management is to restore and enhance species diversity (Newbold et al., 2020), studies are needed to identify the broader effects of different management regimes on wildlife diversity.

Species-rich taxa such as rodents are highly effective systems to measure diversity and infer ecosystem health (Avenant, 2011; Loggins et al., 2019; Fernández et al., 2021). Rodents are the most diverse mammalian taxon, comprising approximately 40% of mammal species worldwide (Burgin et al., 2018), and play crucial roles in ecosystems, including both bottom-up (e.g., seed dispersal; Sunyer et al., 2013) and top-down (i.e., common prey; Geng et al., 2009) processes. Because many rodent species have a fast pace of life strategy (i.e., r-selected), their communities also quickly respond to changes in the environment (Zúñiga et al., 2020). For example, female hispid cotton rats (*Sigmodon hispidus*), a common grassland rodent in the USA, can produce up to 12 pups/litter every four weeks throughout the breeding season (Espinoza and Rowe, 1979).

Despite rodents being an integral part of ecosystems, they also carry many pathogens that can spillover and cause disease in humans (zoonoses; Meerburg et al., 2009; Dahmana et al., 2020). Thus, understanding the impacts of habitat management on pathogens is relevant to both wildlife and human health. Infection dynamics are often shaped by characteristics of individual hosts and their populations. Notably, many pathogens require a minimum host abundance or density to persist in populations (density threshold; Lloyd-Smith et al., 2005) and transmission rates increase as abundance rises (density-dependent transmission; Anderson & May, 1978). In such cases, habitat management could indirectly impact infection dynamics in wildlife and exposure risks for humans by influencing host community diversity and species abundance (Grosholz, 1993; Suzán et al., 2013; Hite et al., 2016).

Research investigating the impacts of habitat variation on the ecology of zoonotic pathogens has primarily focused on a small number of systems with well-established human health implications (e.g., *Peromyscus*-*Borrelia burgdorferi*; Prusinski et al., 2006; Adalsteinsson et al., 2018), but most zoonotic systems are still poorly understood. Drawing meaningful conclusions from pathogen data in wildlife, including rodents, can be difficult due to low prevalence that fluctuates over time and space, and many pathogens are limited to one or a few host species within a community (e.g., Cantoni et al., 2002; Salazar-Bravo et al., 2004; Essbauer et al., 2009). As a result, broad inferences are often made based on model systems rather than specific host-pathogen ecology. For example, most information on American orthohantaviruses is inferred from studies on a select few common viruses despite there being 21 known orthohantaviruses throughout North and South America (Mull et al., 2020).

In this study, we investigate how habitat management impacts wildlife and the viruses they carry. Rodent communities were compared among prairie sites under different management regimes. We assessed the diversity and abundance of rodent species and how this translates to the presence and prevalence (through serology) of three groups of common rodent-borne viruses: orthohantaviruses, arenaviruses, and orthopoxviruses (Forbes et al., 2014; Ogola et al., 2021). Since habitat management is usually designed to enhance species diversity, we hypothesize that rodent diversity and overall abundance will be higher in habitats with management reminiscent of natural ecosystems. We hypothesize the opposite pattern for virus prevalence, with prevalence being lowest in managed habitats, as wildlife hosts of zoonotic pathogens tend to be more common in disturbed habitats (Keesing & Ostfeld, 2021).

## Methods and Materials

### Study sites

Rodents were captured in prairies throughout Benton and Washington Counties, Arkansas, USA. This area lies near the edge of the historical tallgrass prairie ecoregion, and like other tallgrass prairie ecosystems, most of the landscape has been altered by humans, with few remnant prairies remaining. Trapping was conducted at 13 sites within six distinct prairies (Fig. 1). Neighboring sites within the same prairie were considered separate areas due to distinct management regimes and physical barriers (i.e., roadway, riparian habitat, and/or firebreak). Although these barriers do not act as complete physical barriers, they limit rodent movement among sites and distinguish separately-managed parcels. Prairies ranged in size from 6.7-32.6 ha, and distinct management sites within each prairie ranged in size from 1.5-23.6 ha. Site management was classified as one of three regimes: prescribed burning, reminiscent of natural ecosystem functioning (designated burn; five sites); haying, mowing, or other means of mechanical cutting (designated cut; six sites); or no active management of vegetation, leading to heavy woody encroachment (designated control; 2 sites; Fig. 1). Management regimes at the study sites have been continuous for several decades, and our results thus represent long-term effects of management regimes.

**Figure 1.**
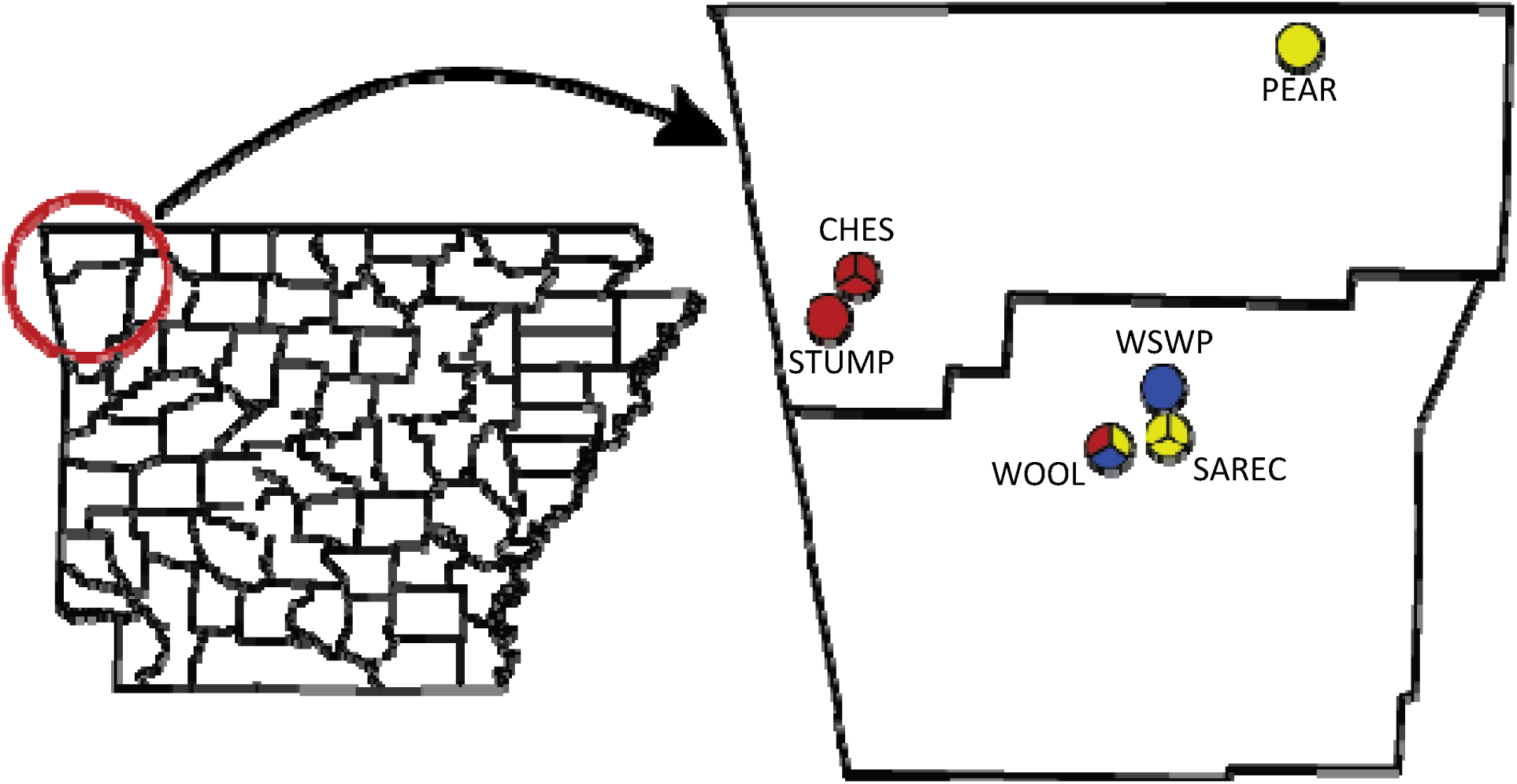
Map of prairie study sites in Benton and Washington Counties, Arkansas, USA. Wedges in circles represent individual sites at each prairie; red wedges indicate burned sites, yellow wedges indicate cut sites, and blue wedges indicate control sites. Each prairie site was given a short name for identification; CHES=Chesney Prairie Natural Area; STUMP=Stump’s Prairie; PEAR=Pea Ridge National Military Park; SAREC=Milo J. Shult Agricultural Research & Extension Center; WOOL=Woolsey Wet Prairie; WSWP=Wilson Springs Wetland Preserve.

### Rodent trapping and sampling

Rodent trapping was conducted once every two months at each site from June-November 2020 and April-July 2021. For each trapping occasion, approximately 50 Sherman live traps (H. B. Sherman Traps, Inc.) were set for two consecutive nights approximately 10m apart in a series of transect lines (see Table 1 for deviations of trap numbers). Traps were baited with a mixture of millet and black oil sunflower seeds and set at dusk. Traps were checked and captured rodents were processed the following morning. Initially, all rodents were euthanized for tissue collection except for species classified as species of conservation need by Arkansas Game and Fish Commission (*Reithrodontomys humulis*, *megalotis*, and *montanus*). Due to permit limitations, we were unable to euthanize all individuals of abundant species (*Reithrodontomoys fulvescens* and *Sigmodon hispidus*) in fall 2020.

**Table 1.** Combined trapping effort and capture success among prairies from 2020-2021. The number of captured, seropositive animals, and Shannon index for all rodents trapped for the duration of the study at each site, with percentages based on the number of trap nights for captures and number of captures for seropositive columns.

| Site | Management | Trap nights | Captures (%) | Seropositive (%) | Shannon index (H) |
| --- | --- | --- | --- | --- | --- |
| CHES_A | Burn | 458 | 60 (13.1) | 8 (13.3) | 1.14 |
| CHES_B | Burn | 458 | 70 (15.3) | 2 (2.9) | 1.18 |
| CHES_C | Burn | 438 | 66 (15.1) | 10 (15.2) | 1.16 |
| STUMP | Burn | 519 | 79 (15.22) | 15 (19.0) | 0.67 |
| WOOL_A | Burn | 500 | 73 (14.6) | 1 (1.4) | 1.06 |
| PEAR_A | Cut | 500 | 11 (2.2) | 0 | 0.86 |
| PEAR_B | Cut | 500 | 7 (1.4) | 0 | 0.60 |
| SAREC_A | Cut | 400 | 20 (5) | 0 | 0.33 |
| SAREC_B | Cut | 400 | 24 (6) | 0 | 0.17 |
| SAREC_C | Cut | 400 | 75 (18.8) | 0 | 0.50 |
| WOOL_B | Cut | 460 | 17 (3.7) | 2 (11.8) | 1.15 |
| WOOL_C | Control | 400 | 14 (3.5) | 0 | 1.33 |
| WSWP | Control | 520 | 100 (19.2) | 0 | 1.33 |
| Total |  | 5953 | 616 (10.3) | 38 (6.2) |  |

Captured rodents were identified to species level based on morphology (pelage and lengths of ear, tail, head/body, and hind foot). Visual inspection was used to determine sex and reproductive condition; males were considered to be reproductive if their testes were descended into the scrotum, and females were considered to be reproductive if their nipples were enlarged or lactating or if their vagina was perforate or plugged. Rodent blood samples were collected via either the submandibular vein directly into a microcentrifuge tube during processing and immediately placed on ice or a heart sample placed into phosphate-buffered saline (PBS) during dissection (see below; Forbes et al., 2014). To promote efficiency and minimize handling time and associated distress to wild rodents, most rodents were quickly euthanized via cervical dislocation without anesthetic; cotton rats were the exception due to their larger size and were anesthetized with inhalation isoflurane prior to cervical dislocation. Euthanized rodents were placed in individual labeled grip-lock bags and stored in a cooler with ice while in the field. Rodents that were not euthanized were ear-tagged and released at their point of capture following sample and data collection.

Euthanized rodents were stored in a -20°C freezer and later dissected under a biosafety hood. Tissue samples were collected aseptically using clean forceps and scissors and placed in sterilized microcentrifuge tubes. Hearts were placed in PBS solution to permit serology assays. All samples and specimens were stored at -20°C.

### Assays to detect antibodies against rodent viruses

Blood samples were tested for antibodies reactive to orthohantaviruses, arenaviruses, and orthopoxviruses using immunofluorescence assays (IFAs), as previously described (Kallio-Kokko et al., 2006; Kinnunen et al., 2011; Forbes et al., 2014). Briefly, samples were diluted in PBS and then incubated on slides with viral antigens followed by several wash cycles to remove unbound antibodies. Fluorescent polyclonal rabbit anti-mouse FITC conjugate was then added to the slides, which were again incubated and washed. Slides were examined under a fluorescence microscope for reactive antibodies. These serology assays are cross-reactive within broad virus groups and therefore are effective and efficient approaches for our non-specific screening (e.g., Ogola et al., 2021).

### Data analyses

All analyses were conducted in R 4.1.0 (R Core Team 2021). We used a chi-square test of independence to compare trapping success among management types. Analysis of variance (ANOVA) and Tukey post-hoc tests were then used to compare Shannon diversity index (H) per site among management regimes to determine the impacts of habitat management on rodent diversity. H values were calculated using the combination of all trapping periods per site to account for seasonal variation in each species. Additionally, to understand how rodent community assemblage varied among management regimes, a chi-square test of independence was used to compare the proportions of grassland specialists (*Microtus ochrogaster*, *Reithrodontomys* spp., and *S. hispidus*) to non-grassland specialist species (*Microtus pinetorum*, *Mus musculus*, and *Peromyscus* spp.).

Because rodents with antibodies against the focus virus groups were only detected in sites that were burned or cut, a chi-square test of independence was used to test for differences in total seroprevalence of all three viruses among habitat management regimes. Binomial generalized linear mixed models (GLMMs) were then used to compare seroprevalence between burned and cut sites, with prairie and site identity as a nested random effect. Demographic data, including sex, reproductive status, abundance index (capture success), and their interactions were set as explanatory variables in the GLMMs. Because most seropositive cases were from rodents with antibodies against orthohantaviruses, we also used GLMMs to compare orthohantavirus seroprevalence alone among burned and cut sites. Two separate binomial GLMMs were used to analyze orthohantavirus seroprevalence from all sites within cotton rats and prairie voles (*M. ochrogaster*), as these two species accounted for the majority of seropositive rodents but have different life histories, including seasonal dynamics and mass (Brady and Slade, 2001). Explanatory variables for rodent species-level GLMMs were mass, sex, reproductive status, abundance index, and their interactions. Finally, a Poisson GLMM was used to compare seroprevalence by trap success at each site and trapping occasion, with trapping occasion as a random effect and prairie and site identity as a nested random effect. GLMMs were conducted using the *lme4* package (Bates et al., 2022); all other statistical analyses were conducted using base R (R Core Team, 2021).

## Results

A total of 616 rodents were captured across 5953 trap nights (Table 1), and no tagged animals were recaptured. Capture success ranged from 0-45% depending on site and season. We captured eight different rodent species throughout the study, with 2-6 different species at individual sites (Fig. 2a).

**Figure 2.**
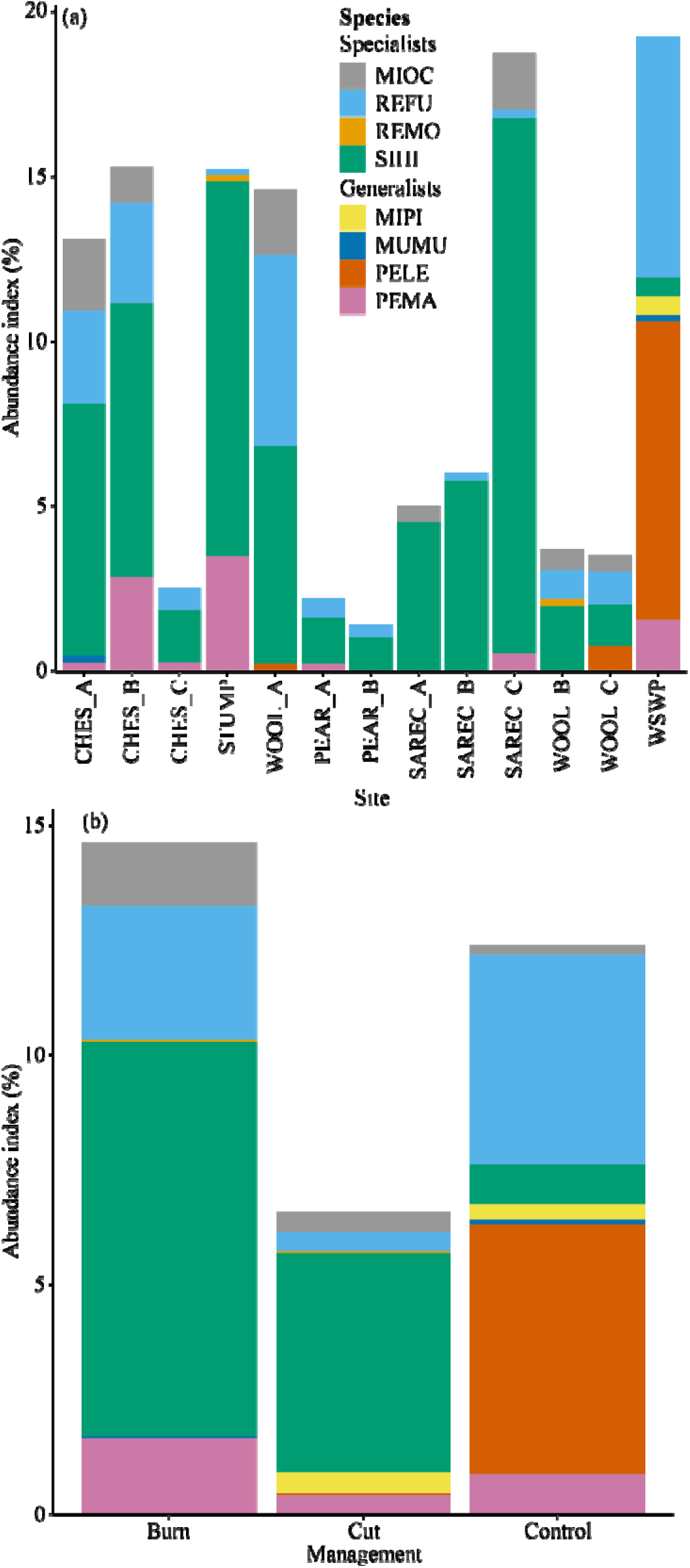
Abundance index of each species per (a) site and (b) management regime. Abundance index is calculated as % capture rate for the duration of the study. Management regime for individual sites can be found in Table 1. Species identifiers are MIOC=*Microtus ochrogaster*; MIPI=*Mi. pinetorum*; MUMU=*Mus musculus*; PELE=*Peromyscus leucopus*; PEMA=*P. maniculatus*; REFU=*Reithrodontomys fulvescens*; REMO=*R. montanus*; SIHI=*Sigmodon hispidus*.

Capture success varied among management regimes (χ*^2^*=91.07, *p*<0.001; Table 1). Success was higher at burned and control sites than cut sites (both *p*<0.001; Fig. 2b) and did not differ between burned and control sites (*p*=0.16). Rodent diversity (H) also varied among habitat management regimes (*F*_(2,10)_=5.18, *p*<0.03; Fig. 3; Table 1), with control (adjusted *p* < 0.05) and burned (adjusted *p*=0.09) sites having higher rodent diversity than cut sites. Rodent diversity did not differ between burned and control sites (adjusted *p*=0.58). Rodent community assemblages also varied among management regimes (χ*^2^*=142.97, *p*<0.001); cut sites had the highest proportion of grassland specialist rodent species, follow by burned and then control sites (all *p*<0.001; Fig. 2).

**Figure 3.**
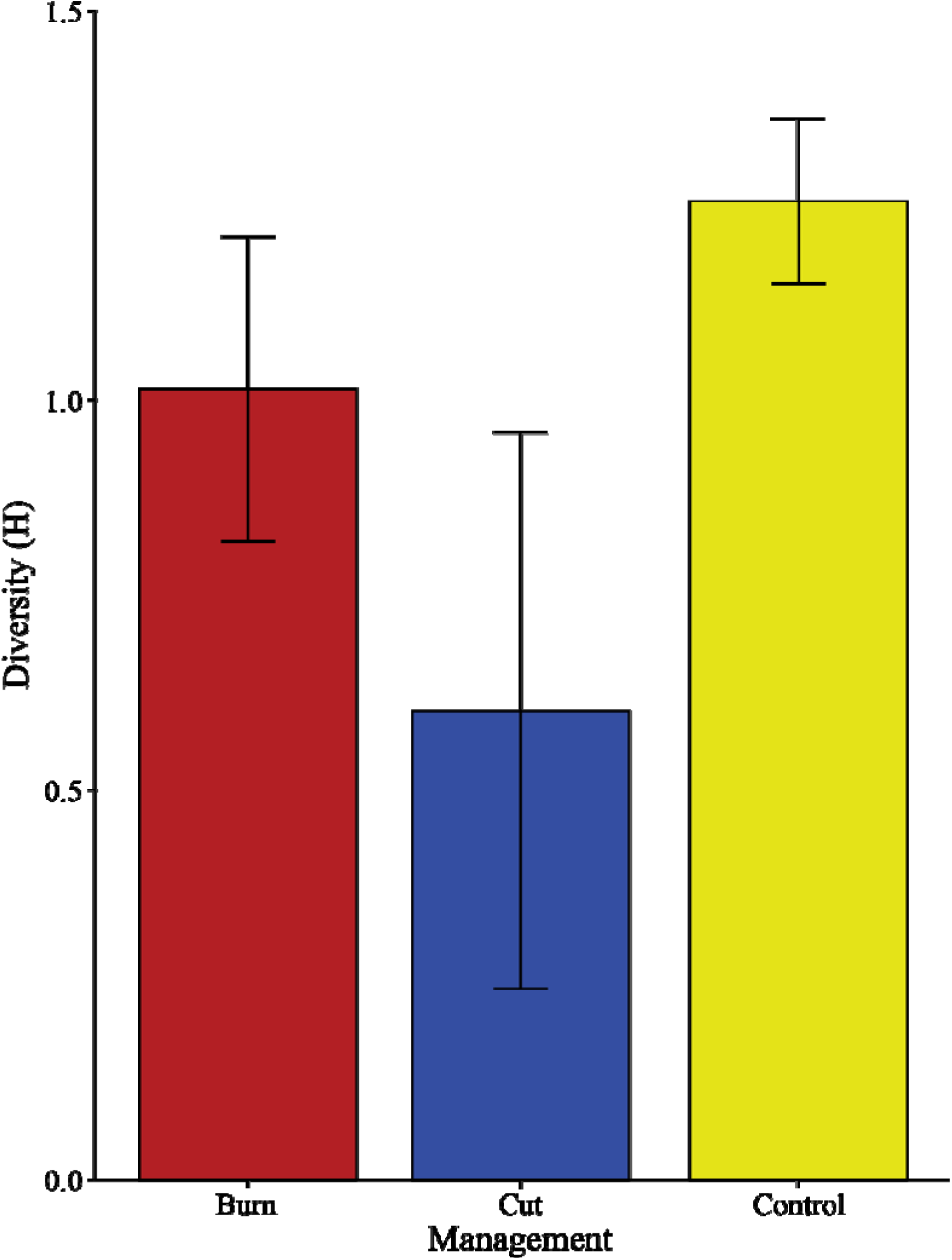
Shannon diversity index (H) among prairies with burned, cut, or control management regimes. Error bars represent ± 1 standard deviation. A single H index is calculated from diversity for the duration of the study per site.

A total of 38 rodents (6.2%) were seropositive for one of the tested virus groups (34 orthohantavirus, 3 arenavirus, 1 orthopoxvirus; Table 2). All seropositive animals were caught at burned sites except one orthohantavirus-seropositive fulvous harvest mouse and one orthohantavirus-seropositive cotton rat (Table 1). The majority of seropositive individuals were cotton rats and prairie voles with orthohantavirus antibodies (Table 2).

**Table 2.**
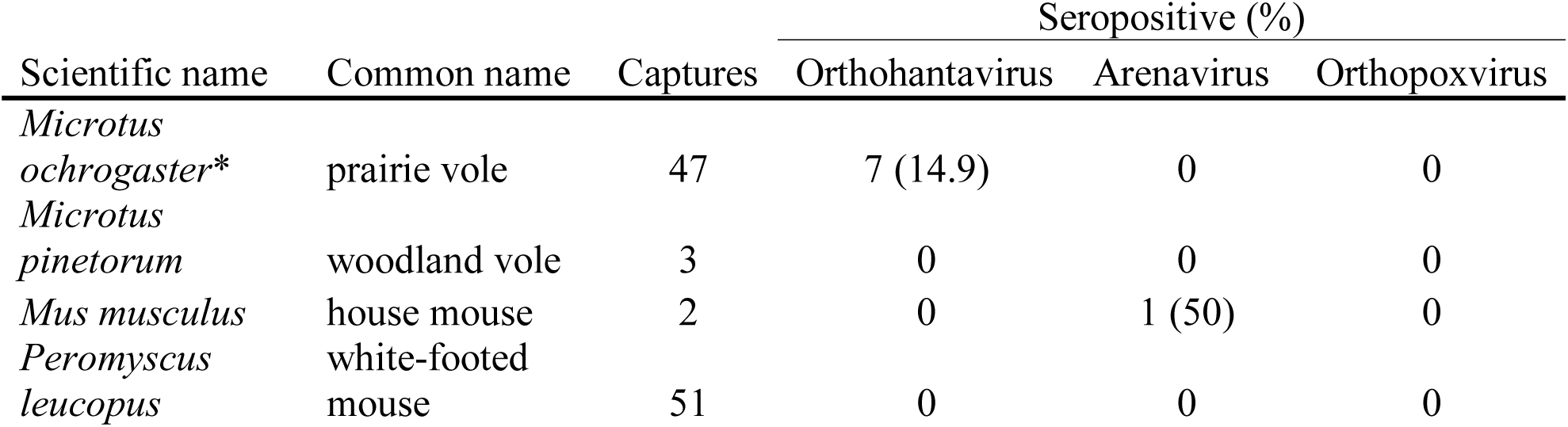
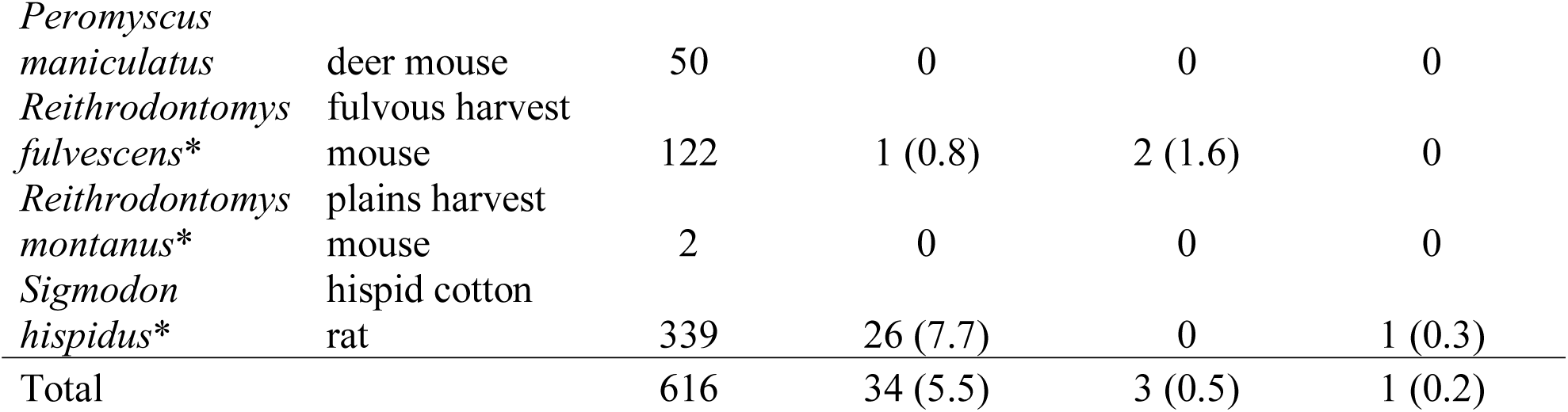
Total number of each species captured among all sites that were seropositive for orthohantavirus, arenavirus, and orthopoxvirus. Grassland specialist species are marked with an *.

Complete processing data was collected for 609/616 rodents captured for infection analyses. Based on the Chi-square test for independence, virus seroprevalence varied among management types (χ^2^=24.69, *p*<0.001; Fig. 4), with a higher proportion of seropositive rodents in burned sites than cut or control sites (both *p*<0.001). No difference in seroprevalence was detected between cut and control sites (*p*=0.61). The most parsimonious GLMM comparison between burned and cut sites only included type of habitat management and reproductive condition (Table S1). This model confirmed that seropositive rodents were more common in burned sites than cut sites (*p*<0.04) and that reproductive individuals were more likely to be seropositive than non-reproductive individuals (*p*<0.01). Unsurprisingly, orthohantavirus seroprevalence was similar to overall seroprevalence, with burned sites and reproductive condition being predictors of orthohantavirus seropositivity (*p*<0.05 and *p*<0.01, respectively). Additionally, higher rodent abundance was associated with greater seroprevalence at a given site and trapping occasion (*p*<0.03).

**Figure 4.**
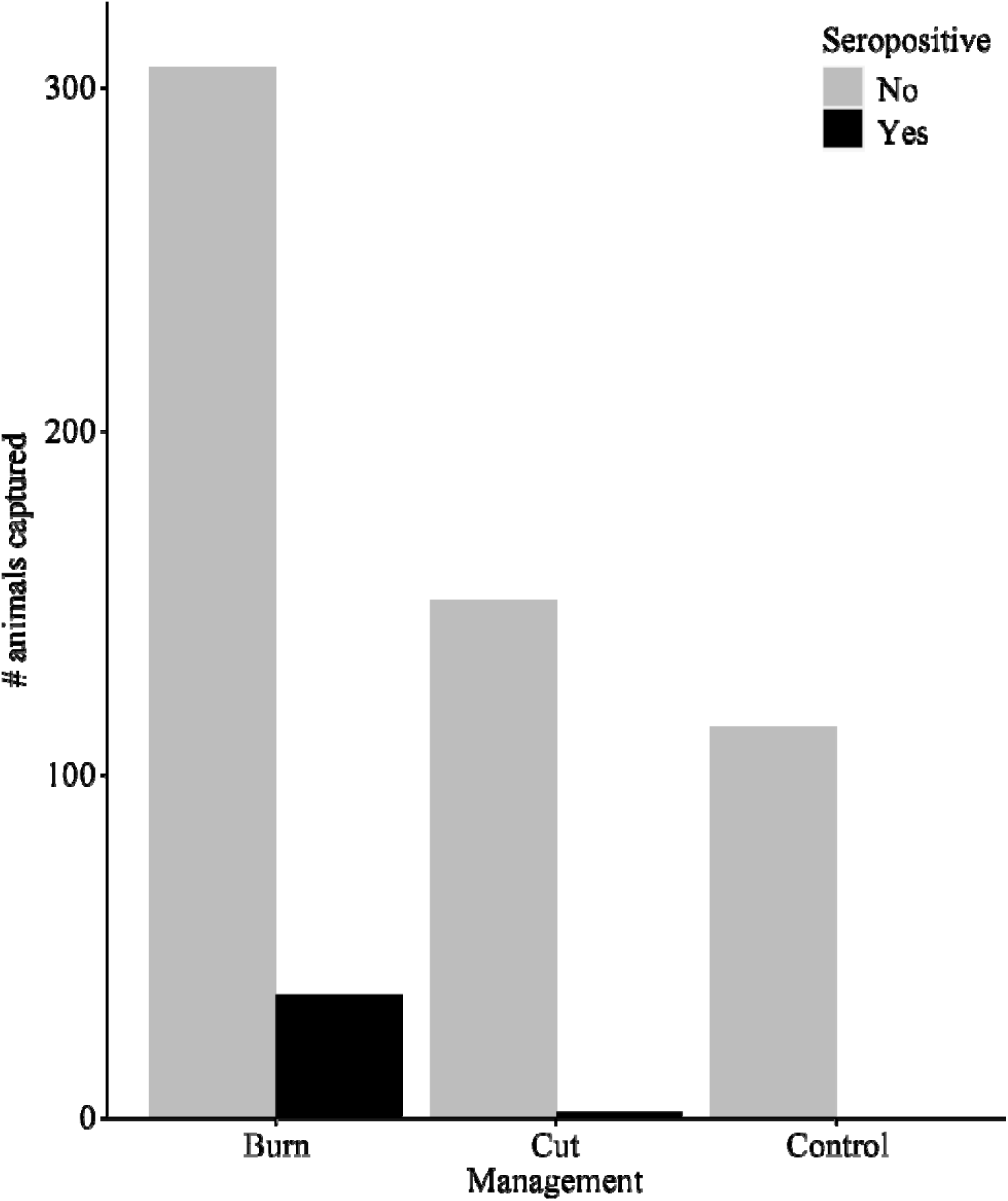
Number of rodents caught among sites with burned, cut, or control management regimes for the duration of the study that were seropositive or seronegative for any tested virus group (orthohantaviruses, arenaviruses, or orthopoxviruses).

Similar to the GLMM with all individuals, sex and abundance were not important predictors in seroprevalence for cotton rats or prairie voles (Tables S2 and S3). However, reproductive condition was not a variable in the most parsimonious models at the species level. Heavier individuals of both cotton rats and prairie voles were more likely to be seropositive than lighter individuals (*p*<0.001 and *p*<0.04, respectively; Fig. 5).

**Figure 5.**
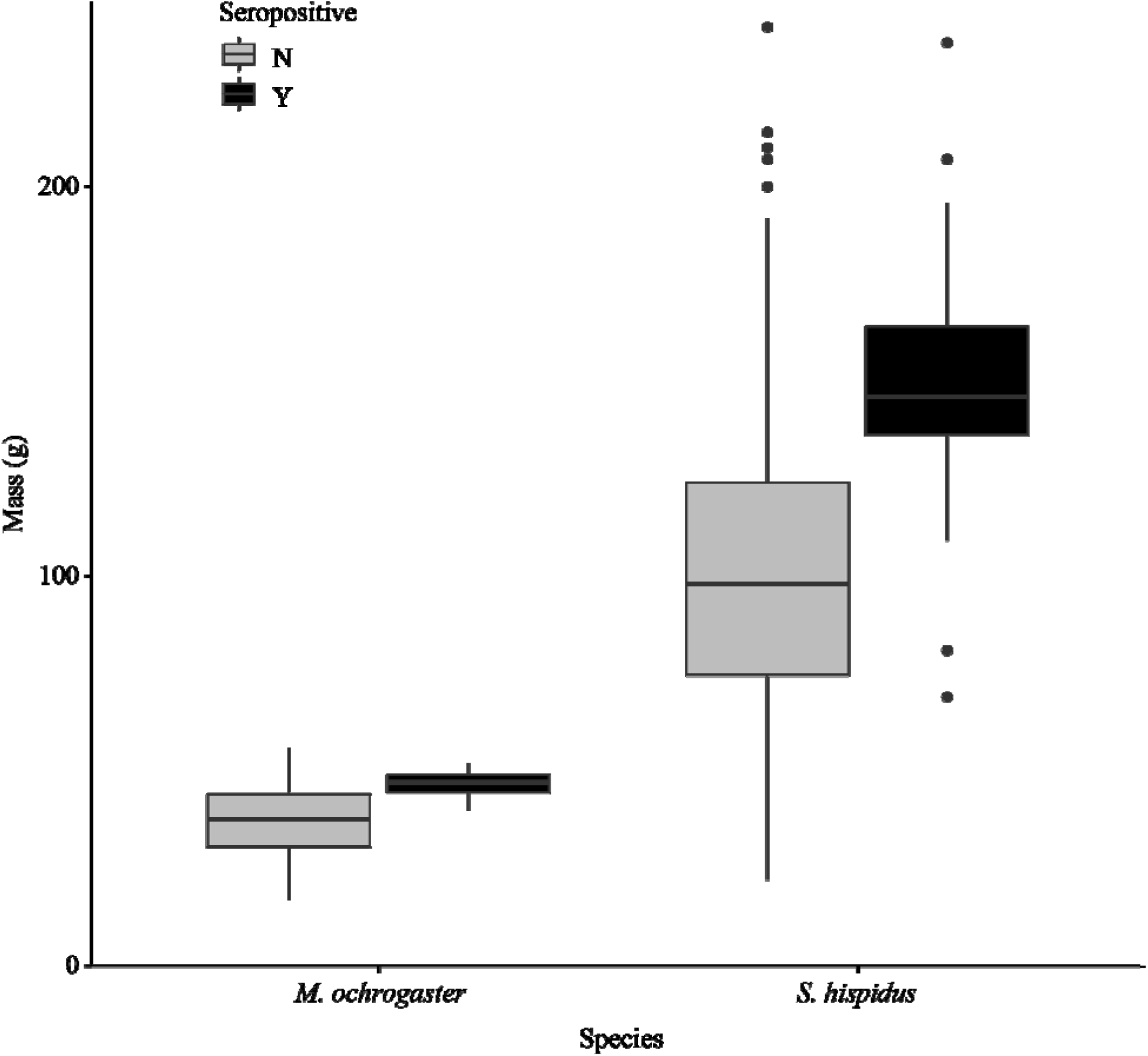
Mass of *Microtus ochrogaster* (prairie voles) and *Sigmodon hispidus* (hispid cotton rats) from all sites that were seropositive or seronegative for orthohantaviruses.

## Discussion

We demonstrate that habitat management regimes lead to differences in rodent community assemblages, species abundances, and subsequently, viral infection dynamics. Burned habitats produced the highest overall quality of grassland rodent communities, with high rodent diversity, overall abundance, and a relatively high proportion of grassland specialist species. In comparison, cut habitats had low diversity and abundance but a high proportion of grassland specialists, and control habitats had high diversity and abundance but a low proportion of grassland specialists. However, most of the virus seropositive rodents in this study were grassland specialists found in burned sites. Our findings highlight the advantages and disadvantages of different grassland habitat management regimes for biodiversity indicators and the importance in identifying these tradeoffs.

Although sites were grouped according to management regime, some differences in management schedules, site history, and biogeochemical factors were unavoidable and created heterogeneity within group categories. In particular, three of the five burned sites were burned every three years and the other two were burned annually. These differences were unavoidable due to the study design, akin to a natural experiment. However, potential differences due to site heterogeneity within management categories were assessed to validate groupings; no differences in rodent abundance, rodent diversity, or seroprevalence were detected between burn frequencies (Appendix S1). Despite several replicates of burned and cut sites, only two control sites were available in this study, as these habitats change drastically with the onset of management and are prone to ecological succession in the prolonged absence of management.

We found that burned and control sites had similar rodent diversity and overall abundance but differed in the relative proportions of grassland specialists. The high proportion of generalist and forest-specialist species in control sites is indicative of the consequences of habitat degradation in the absence of prairie management, particularly from woody encroachment and loss of non-woody diversity (Miller et al., 2000; Brunsell et al., 2017). Although some studies have shown that burning increases the relative abundance of habitat generalists (e.g., Manyonyi et al., 2020; Zúñiga et al., 2020), such outcomes generally represent the immediate effects of fire, as opposed to prolonged effects from a decade or more of management. Indeed, rodent diversity at several of our burned sites varied considerably from a previous assessment shortly after active management began (Nelson, 2005), most notably by an increase in our study of prairie voles and fulvous harvest mice, two grassland specialist species (Table 2). Similar positive long-term effects of prescribed burning on habitat availability and species richness in grasslands have been identified for other wildlife taxa, including insects and elk (Van Dyke & Darragh, 2006; Bargmann et al., 2015; Podgaiski et al., 2017).

Cut sites, on the other hand, had lower diversity and abundance but a higher relative proportion of grassland specialists than burned and control sites. Hayed or mowed fields generally have low vegetation diversity compared to other grasslands (Faria et al., 2018), which limits ecosystem functioning across a variety of wildlife (Wan et al., 2020). Species capable of using the dominant vegetation in these areas have access to abundant resources and their population sizes can become very large. For example, one of our study cut sites had the second highest abundance, and 87% of the rodents captured were hispid cotton rats (Table 1). Grazing by livestock is another method to manage habitats that is often considered analogous to cutting vegetation. Although grazing is a more natural means of removing vegetation and can produce more diverse vegetation communities than haying or mowing (Tälle et al., 2016), intensive grazing drastically reduces rodent abundance (Yarnell et al., 2007; La Morgia et al., 2015). Thus, while light grazing may increase wildlife diversity, intensive grazing is likely similar to, or worse than, cutting for rodent diversity.

The high diversity and abundance of grassland specialists in burned sites is likely responsible for the differences in virus seroprevalence among habitat management regimes, as all seropositive animals were grassland specialists. In particular, cotton rats, the reservoir host of Black Creek Canal virus (orthohantavirus; Rollin et al., 1995), were the most commonly trapped species and accounted for the majority of seropositive individuals for all viruses and orthohantaviruses specifically (Tables 2 and S4). High virus prevalence in dominant rodent species in natural, pristine habitats is consistent with recent studies from South America (Burgos et al., 2021; Tirera et al., 2021). Interestingly, we identified no seropositive deer mice or white-footed mice, the reservoir hosts of Sin Nombre virus and New York virus (both orthohantaviruses that would be detected with our serology assay; Yamada et al., 1994), respectively (Childs et al., 1994; Hjelle et al., 1995), despite relatively high orthohantavirus seroprevalence in the similarly-abundant prairie voles (Table 2). Thus, given that seroprevalence was dominated by orthohantaviruses in grassland specialists, the low density of grassland species in control sites, and of all rodent species in cut sites, was likely insufficient for viral establishment and persistence in these habitats (Mills et al., 1999).

In addition to management regime, several other variables influenced infection dynamics in this study. Although rodent abundance was important for seroprevalence at the site level, it was not important at the individual level. This is likely due to a time lag effect, where prevalence is impacted by earlier rather than current density (Yates et al., 2002; Adler et al., 2008). Heavier individuals, which was our correlate for age class, and to an extent body condition, were more likely to be seropositive for orthohantavirus (Fig. 5), consistent with previous studies (Glass et al., 1998; Walsh et al., 2007). There were too few rodents seropositive for arenaviruses or orthopoxviruses for statistical evaluation, but the high demographic variety of arenavirus hosts in this study (Table S4) corroborates our understanding of American arenavirus host demography, which includes individuals of both sexes and all age classes (Milazzo et al., 2008; Milazzo et al., 2013). Conversely, the low seroprevalence of orthopoxviruses was surprising, as these viruses are commonly found in diverse wild rodent species from other geographical areas (e.g., Kinnunen et al., 2011; Forbes et al., 2014; Ogola et al., 2021), though relatively little is known about orthopoxviruses in American rodents (but see Emerson et al., 2009).

Overall, our study evaluates the impacts of long-term habitat management on wildlife and their pathogens. As expected, high intensity grassland management (i.e., prescribed burning) generated high diversity and abundance of rodents. Burned sites also had the highest virus seroprevalence and were the only sites where rodents with antibodies against two of the three virus groups were detected. Biodiversity is crucial for healthy ecosystem functioning, and these results provide empirical evidence that can inform grassland prairie restoration and ongoing management strategies.

## Acknowledgements

We thank Reilly Jackson, Ellery Lassiter, Shannon Kitchen, Aaron Norris, C. Houston Lamb, and Alexia Hernandez for assistance with rodent trapping. We also thank Sanna Mäki and other staff of the Viral Zoonosis Research Unit from the University of Helsinki for assistance with serology tests. We are very grateful to the prairie land managers for allowing us to collect samples from and for managing the sites used in this study, including Arkansas Natural Heritage Commission, City of Fayetteville, Joe Woolbright, Milo J Shult Agricultural Research & Extension Center, Northwest Arkansas Land Trust, and Pea Ridge National Military Park. This study was conducted under the University of Arkansas Institutional Animal Care and Use Committee (IACUC) permit number 20028 and Arkansas Game and Fish Commission permit numbers 102820194 and 030820211. This research was partly supported by NSF DEB 1911925.

## Authors’ Contributions

The initial ideas were conceived by N.M. and K.M.F.; N.M., A.Sc., A.St., and T.S. collected the data; K.M.F. provided resources; N.M. analyzed the data; N.M. drafted the initial manuscript with input from all authors. All authors contributed to draft revisions and approved the final version for publication.

## Data Accessibility

All data will be made publicly available on Dryad upon acceptance of the manuscript for publication.

