## Supplemental Appendix 1 for "Effects of habitat management on rodent diversity, abundance, and virus infection dynamics"

**Appendix S1.** As a form of validation, we compared capture success, rodent diversity, and rodent seroprevalence between sites that were burned every three years and sites that were burned annually to verify that burn frequency did not influence our results. Capture success was compared using a Chi-square test for independence (χ^2^=0.063, *p*=0.80). Rodent diversity was compared using a linear mixed effects model (*lme4* package in R) with each site’s Shannon index as the response variable, burn frequency as the explanatory variable, and prairie as a random effect (*p*=0.60). Rodent seroprevalence was compared using a binomial generalized linear mixed model (*lme4* package in R) with individual seroprevalence as the explanatory variable, burn frequency as the explanatory variable, and prairie as a random effect (*p*=0.89).
