## Supplemental Table 1 for "Effects of habitat management on rodent diversity, abundance, and virus infection dynamics"

**Table S1. AIC values for GLMMs comparing seroprevalence of all rodents between burned and cut sites.**

| Variables | AIC |
| --- | --- |
| Management * Sex | 252.2 |
| Management * Success + Sex | 251.9 |
| Management * Sex + Success | 251.9 |
| Management * Success | 250.7 |
| Management + Sex | 250.5 |
| Management + Success + Sex | 250.2 |
| Management | 249.1 |
| Management + Success | 249.0 |
| Management * Sex + Success + Reproductive | 244.8 |
| Management * Success + Sex + Reproductive | 244.4 |
| Management * Reproductive + Success + Sex | 244.4 |
| Management * Sex + Reproductive | 243.2 |
| Management + Success + Sex + Reproductive | 243.0 |
| Management * Reproductive + Sex | 242.9 |
| Management * Reproductive + Success | 242.8 |
| Management + Sex * Reproductive | 242.8 |
| Management * Success + Reproductive | 242.7 |
| Management + Sex + Reproductive | 241.4 |
| Management * Reproductive | 241.2 |
| Management + Reproductive | 239.8 |
