## Supplemental Table 2 for "Effects of habitat management on rodent diversity, abundance, and virus infection dynamics"

**Table S2. AIC values for GLMMs comparing orthohantavirus seroprevalence of hispid cotton rats (*Sigmodon hispidus*).**

| Variables | AIC |
| --- | --- |
| Mass * Sex * Reproductive * Abundance | 147.2 |
| Mass * Sex * Abundance + Reproductive | 143.8 |
| Mass * Abundance * Reproductive + Sex | 140.7 |
| Mass * Sex * Reproductive + Abundance | 140.7 |
| Mass * Sex * Reproductive | 140.3 |
| Mass * Sex + Reproductive + Abundance | 139.4 |
| Mass * Sex + Reproductive | 138.9 |
| Mass + Sex + Reproductive * Abundance | 138.1 |
| Mass * Abundance + Sex + Reproductive | 138.0 |
| Mass + Sex * Reproductive + Abundance | 137.8 |
| Mass + Sex * Abundance | 137.5 |
| Mass + Sex + Reproductive + Abundance | 137.5 |
| Mass * Sex + Abundance | 137.4 |
| Mass * Reproductive + Sex + Abundance | 137.4 |
| Mass + Sex * Reproductive | 137.3 |
| Mass + Reproductive * Sex | 137.3 |
| Mass + Sex + Reproductive | 137.0 |
| Mass * Reproductive + Sex | 136.9 |
| Mass * Abundance + Reproductive | 136.1 |
| Mass * Abundance + Sex | 136.0 |
| Mass + Reproductive + Abundance | 135.6 |
| Mass * Reproductive + Abundance | 135.6 |
| Mass + Sex + Abundance | 135.5 |
| Mass * Reproductive | 135.0 |
| Mass + Reproductive | 135.0 |
| Mass * Abundance | 134.2 |
| Mass + Abundance | 133.7 |
| Mass | 133.3 |
