## Supplemental Table 3 for "Effects of habitat management on rodent diversity, abundance, and virus infection dynamics"

**Table S3. AIC values for GLMMs comparing seroprevalence of prairie voles (*Microtus ochrogaster*). Interaction effects between reproductive and other variables could not be computed because all seropositive prairie voles were in reproductive condition.**

| Variables | AIC |
| --- | --- |
| Mass + Sex * Abundance + Reproductive | 35.4 |
| Mass * Abundance + Sex + Reproductive | 34.5 |
| Mass * Sex + Abundance + Reproductive | 34.1 |
| Mass + Sex * Abundance | 34.0 |
| Mass + Sex + Abundance + Reproductive | 33.8 |
| Mass * Abundance + Sex | 33.7 |
| Mass * Abundance + Reproductive | 33.2 |
| Mass + Sex + Reproductive | 33.2 |
| Mass * Abundance | 33.2 |
| Mass * Sex + Reproductive | 32.9 |
| Mass + Sex + Abundance | 32.7 |
| Mass + Success + Reproductive | 32.5 |
| Mass + Sex | 32.5 |
| Mass + Abundance | 31.8 |
| Mass + Reproductive | 31.5 |
| Mass * Sex | 31.3 |
| Mass | 31.3 |
