## Supplemental Table 4 for "Effects of habitat management on rodent diversity, abundance, and virus infection dynamics"

**Table S4. Individual demographic information for seropositive rodents. For virus, H=orthohantavirus, A=arenavirus, and P=orthopoxvirus.**

| Virus | Species | Sex | Reproductive | Mass (g) | Site |
| --- | --- | --- | --- | --- | --- |
| H | *Microtus ochrogaster* | F | Yes | 47 | CHES_C |
| H | *Microtus ochrogaster* | F | Yes | 48 | CHES_C |
| H | *Microtus ochrogaster* | F | Yes | 52 | CHES_A |
| H | *Microtus ochrogaster* | M | Yes | 40 | CHES_A |
| H | *Microtus ochrogaster* | M | Yes | 44 | CHES_C |
| H | *Microtus ochrogaster* | M | Yes | 45 | CHES_B |
| H | *Microtus ochrogaster* | M | Yes | 50 | CHES_A |
| H | *Reithrodontomys fulvescens* | M | Yes | 11 | WOOL_A |
| H | *Sigmodon hispidus* | F | No | 126 | STUMP |
| H | *Sigmodon hispidus* | F | No | 133 | CHES_C |
| H | *Sigmodon hispidus* | F | No | 136 | STUMP |
| H | *Sigmodon hispidus* | F | Yes | 81 | WOOL_B |
| H | *Sigmodon hispidus* | F | Yes | 155 | STUMP |
| H | *Sigmodon hispidus* | F | Yes | 167 | CHES_C |
| H | *Sigmodon hispidus* | F | Yes | 169 | CHES_A |
| H | *Sigmodon hispidus* | F | Yes | 174 | STUMP |
| H | *Sigmodon hispidus* | M | No | 139 | STUMP |
| H | *Sigmodon hispidus* | M | No | 153 | CHES_C |
| H | *Sigmodon hispidus* | M | No | 207 | CHES_C |
| H | *Sigmodon hispidus* | M | Yes | 69 | CHES_C |
| H | *Sigmodon hispidus* | M | Yes | 109 | STUMP |
| H | *Sigmodon hispidus* | M | Yes | 129 | CHES_C |
| H | *Sigmodon hispidus* | M | Yes | 137 | STUMP |
| H | *Sigmodon hispidus* | M | Yes | 139 | CHES_A |
| H | *Sigmodon hispidus* | M | Yes | 140 | STUMP |
| H | *Sigmodon hispidus* | M | Yes | 142 | CHES_A |
| H | *Sigmodon hispidus* | M | Yes | 146 | WOOL_B |
| H | *Sigmodon hispidus* | M | Yes | 146 | STUMP |
| H | *Sigmodon hispidus* | M | Yes | 148 | CHES_B |
| H | *Sigmodon hispidus* | M | Yes | 159 | STUMP |
| H | *Sigmodon hispidus* | M | Yes | 164 | STUMP |
| H | *Sigmodon hispidus* | M | Yes | 164 | STUMP |
| H | *Sigmodon hispidus* | M | Yes | 196 | STUMP |
| H | *Sigmodon hispidus* | M | Yes | 237 | STUMP |
| A | *Mus musculus* | M | Yes | 19 | CHES_A |
| A | *Reithrodontomys fulvescens* | F | Yes | 16 | CHES_V |
| A | *Reithrodontomys fulvescens* | M | Yes | 9 | STUMP |
| P | *Sigmodon hispidus* | M | Yes | 133 | CHES_A |
